# Life-cycle mediated effects of urbanization on parasite communities in the estuarine fish, *Fundulus heteroclitus*

**DOI:** 10.1101/404756

**Authors:** James M. Alfieri, Tavis K. Anderson

## Abstract

This study examined the relationship between urbanization and parasite community structure in the estuarine fish, *Fundulus heteroclitus*. We measured landscape and physicochemical factors associated with urbanization at 6 sites from 4 collection periods. Concurrently, we quantified the metazoan parasite community in *F. heteroclitus* collected at those sites, with 105 fish studied per site during the 4 collection periods. Parasite community composition differed between sites. Variation in the prevalence and intensity of infection of two indirect life-cycle parasites, *Lasiocotus minutus* and *Glossocercus caribaensis*, were the primary parasite species that determined this pattern. Sediment potassium and aquatic osmium were the most important physicochemical factors in structuring parasite communities, and habitat dominance was the most important landscape factor. Our data supports the hypothesis that urbanization, acting at both landscape and physicochemical scales, can have a significant impact on parasite community structure. This, however, varied by parasite life history: there was little effect of urbanization on the prevalence and intensity of direct life-cycle parasites, but significant variation was dedicated for indirect life-cycle parasites. This study demonstrates how anthropogenically driven landscape change influences fine-scale parasite population dynamics.

## Introduction

Urbanization significantly alters estuarine environments through landscape composition changes, habitat fragmentation, and contamination (Paul and Meyer, 2001; Lowe and Peterson, 2014; Lerberg et al. 2000). These anthropogenic modifications can affect the diversity and population dynamics of free-living animals and their associated parasite communities (Lotze et al., 2006; Hechinger et al. 2007). Individual parasite species may be affected by urbanization through direct exposure to the abiotic environment (e.g. contaminants and altered physicochemical parameters) (Anderson and Sukhdeo, 2010; Blanar et al. 2011) or through changes in host population dynamics (Anderson and Sukhdeo, 2013a). Parasite species co-occurring in a host species – the parasite community – simultaneously use the same host resource, but each parasite species may have a different life history strategy. Notably, indirect life-cycle parasites have life stages that are exposed to the external environment and are dependent upon trophic interactions; whereas direct life-cycle parasite transmission largely does not involve trophic interactions. Consequently, urbanization may differentially affect parasite species but data that associates urbanization with parasite communities and life-history strategy are sparse (but see Calegaro-Marques 2014 and Blanar et al. 2016).

Parasite life-cycles can be broadly characterized as either indirect or direct. Indirect life-cycle parasites require different species of hosts to complete their life-cycle and are largely dependent on trophic interactions which occur more frequently in locally stable communities (Anderson and Sukhdeo, 2013a). This essential interaction has been used to demonstrate how parasites can be indicators of free-living diversity (Hechinger et al., 2007), and as such, it has been argued that a healthy ecosystem is one that is rich in parasites (Hudson et al., 2006). In urbanized estuaries, host population dynamics may be less stable, resulting in reduced diversity or less predictable host dynamics, causing a concomitant decrease in the richness and abundance of indirect life-cycle parasites (Anderson and Sukhdeo, 2013b). On the other hand, direct life-cycle parasites require only one species of host to complete their life-cycle and are largely dependent on host population densities, as abundant hosts increase the probability of a parasite encountering a suitable environment to colonize (Arneberg et al., 1998). A feature of urbanized estuaries is habitat fragmentation leading to small patch sizes, which can result in increased host densities, causing a concomitant increase in the richness and abundance of direct life-cycle parasites (Wang and Moskovits, 2001; Hussain et al. 2013).

Many of the studies that examine the interaction between host and parasite in urbanized ecosystems have been conducted with human pathogens (Bradley and Altizer, 2007), consider a single host and a single parasite (Johnson et al., 2009), or a single contaminant and landscape factor. Further, linking the regional process of urbanization to local transmission and diversity of parasite communities has yet to reveal consistent patterns (Schotthoefer et al., 2011; Chapman et al., 2015; Blanar et al., 2016; Johnson et al., 2016). One potential explanation for the absence of consistent results is that urbanization differentially affects parasite species: the size and direction of effect is contingent upon parasite life history. In this study, we assessed the effect of multiple landscape composition and physicochemical factors on 24 parasite communities of the salt marsh fish, *Fundulus heteroclitus*. The parasite community in *F. heteroclitus* includes indirect and direct life-cycle species, and the impact of urbanization on the structure parasite communities should be mediated through different life-cycle strategies. We predicted that urbanized sites would have low diversity and prevalence of complex life-cycle parasites; but direct life-cycle parasites would increase in diversity and abundance.

## Materials and methods

### Study site description

We examined 6 sites along the coastline of Georgia, USA, that reflected variation in urbanization, measured through % impervious surface within a 250 m radius around each site (Fig 1). We used this metric to select sites because impervious surface is a strong predictor of different effects of urbanization, such as landscape composition changes, habitat fragmentation, and contaminants (Paul and Meyer, 2001; Lerberg et al. 2000). The 250 m radius around each site was chosen because this is similar to the home range size of the focal host organism, *Fundulus heteroclitus* (Lotrich, 1975; Teo and Able, 2003). The 250 m radius criteria was used as an objective measure to select sites representing a range of urbanization and its effects, however, for the landscape composition analyses, we used a larger spatial scale at the watershed level that is described below.

**Fig 1.**
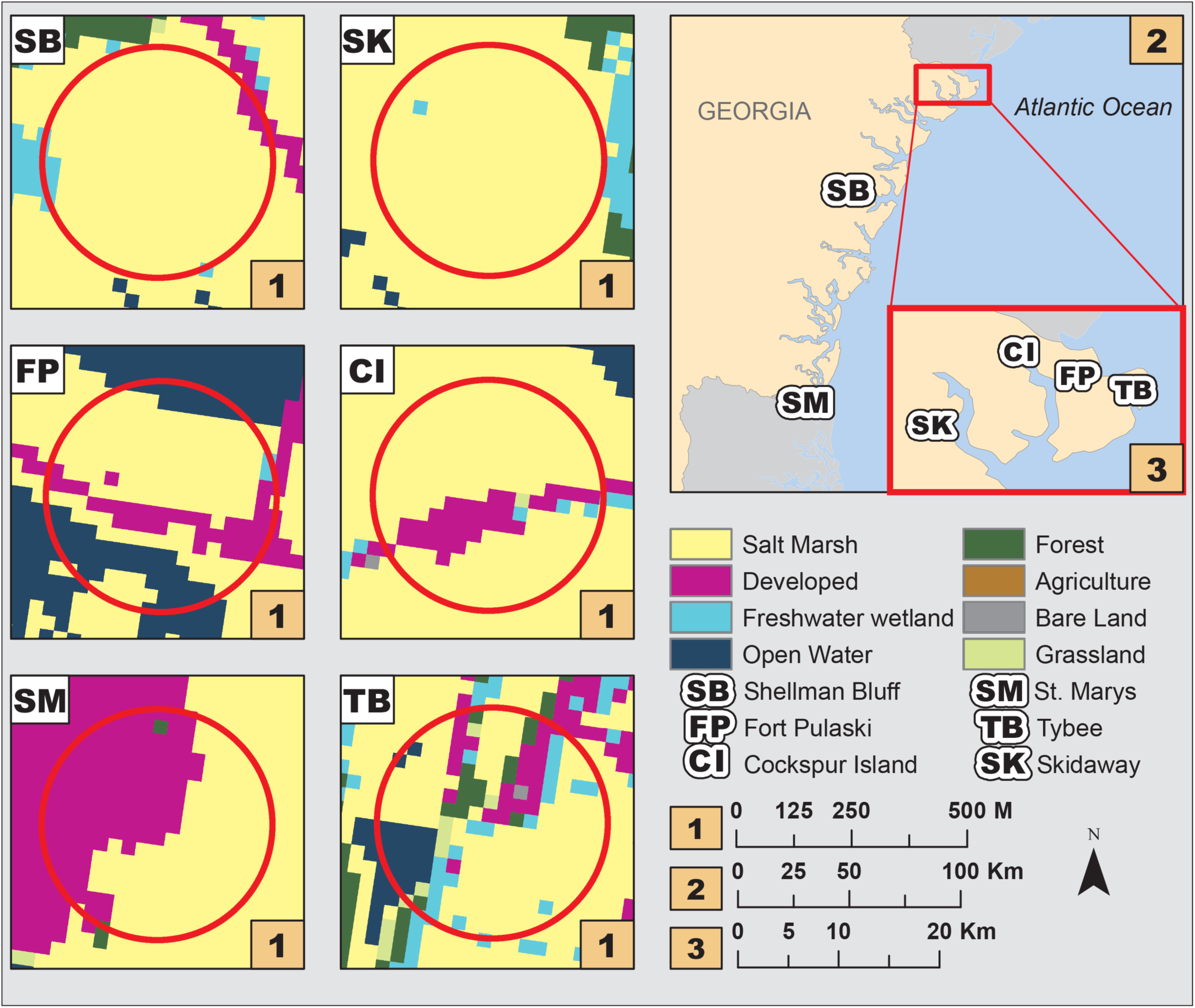
Map of the 6 salt marsh study sites in coastal Georgia, USA, including 2011 land cover data. The sites include Fort Pulaski (FP), Cockspur Island (CI), Tybee (TB), and Skidaway Island (SK) in the Savannah River Estuary; Shellman Bluff (SB) as a representative of the Sapelo Sound; and St. Marys (SM) as part of the St. Marys Estuary. Red circles represent a 250 m buffer around the study site. For the purpose of this work, land cover data was reclassified from the original 21 categories to 10 (Salt Marsh, Developed, Freshwater Wetland, Open Water, Forest, Agriculture, Bare Land, and Grassland): 2 land cover categories were not present in the study area (Freshwater Aquatic Bed and Unconsolidated Shore).

### Quantifying the landscape and physicochemical environment

We defined the boundaries of the landscape to the lowest level watershed provided by the United States Geological Survey (USGS) Watershed Boundary Dataset (WBD): the hydrological unit code (HUC) 12-digit boundary (Lotrich, 1975). We used the 2011 National Land Cover Dataset (NLCD) and National Oceanic and Atmospheric Administration’s (NOAA) Coastal Change Analysis Program (C-CAP) for the regional land cover analysis. Regional landscape composition and change were measured using ArcGIS (ArcGIS Version 10.3, Environmental Systems Research Institute, Redlands, California). We also calculated a series of regional landscape metrics: patch density, largest patch index, edge density, mean salt marsh size, % salt marsh, % developed land, and total area using FRAGSTATS software (FRAGSTATS, version 4.2, University of Massachusetts, Amherst, Massachusetts). Patch density is an index that expresses the number of continuous habitat types, or patches, per unit area, and is a metric for landscape heterogeneity. Largest patch index is the size of the largest habitat patch relative to the size of the landscape, and is a measure of landscape dominance. Edge density is an index of the amount of habitat edges per unit area, and the contrast between developed land and salt marsh was weighted at 0.5 to evaluate salt marsh habitat fragmentation by urban structures or landscapes. Mean salt marsh size is an index of the size of a salt marsh habitat patch, with larger patches weighted in order to decrease the sensitivity of the mean to small habitat patch sizes. The % salt marsh and developed land is the percentage of the landscape comprised of those categories, and is a metric for habitat size and dominance. Total area refers to the area of the watershed. In addition, we used the National Oceanic and Atmospheric Administration’s (NOAA) Coastal Change Analysis Program (C-CAP) to evaluate % landscape change within the HUC-12 digit watersheds for the period of time from 1996 – 2010.

We collected information on the physicochemical environment from each site at flow and ebb tide in March, June, September 2015 and February 2016. Flow and ebb tide measurements were averaged for each site and time point for analysis. We measured pH, temperature, salinity, and conductivity using a handheld multiparamenter instrument (Yellow Springs Instruments Professional Plus; Yellow Springs Instruments, Yellow Springs, Ohio). In addition, we quantified the trace metal composition, nutrient concentrations, and chlorophyll *a* concentration during these dates. To extract chlorophyll *a*, we followed a standard acidification method (EPA 445.0), whereby a volume of 100 ml was filtered using Whatman GF/F filters (47 mm; 0.7 μm nominal pore size). Phytoplankton pigments were then extracted from the filters in 8 ml of 90 % acetone at 0 °C for 24 hours. Fluorescence was measured using a Trilogy fluorometer (Turner Designs, San Jose, California). For the nutrient and trace metal quantification, 2 samples from each collection time and site were sent to the University of Georgia Laboratory for Environmental Analysis (Athens, Georgia). These samples were filtered using a microwave-assisted nitric acid method (EPA 3015), and total nitrogen, total phosphorus, and 72 metals were analyzed with inductively coupled plasma mass spectrometry (ELAN 9000; PerkinElmer, Waltham, Massachusetts). In addition to water analyses, we quantified the trace metal composition of sediment by collecting 3 - 8 cm core samples from randomized locations within each site in September 2015. These samples were sent to the University of Georgia Laboratory for Environmental Analysis (Athens, Georgia), and were prepared using a microwave-assisted nitric acid method (EPA 3052).

### Quantifying the parasite community in *Fundulus heteroclitus*

We used the parasite community of a focal species, the common marsh killifish *Fundulus heteroclitus*, to assess the impact of human modification of the environment on parasite establishment and richness. The host community (benthic invertebrates, birds, fishes) transmits at least 22 species of helminth parasites to the common killifish (Harris and Vogelbein 2006). At each timepoint, 5 baited minnow traps were randomly placed at every site, during the flood tide and collected during the ebb tide. We combined all 5 traps, and randomly subsampled to 30 killifish to obtain a representative population of each site and date combination. The fish were separately housed in the animal facility at Georgia Southern University and necropsied within 1 week (IACUC #I14013).

Each fish was euthanized in a buffered 300 mg L^-1^ solution of tricaine methanesulfonate (MS-222) until cessation of opercula movement followed by sex determination and measurements of total length and weight. The MS-222 solution, all external surfaces, gills, and opercula were examined for ectoparasites and helminths. The viscera, gonads, spleen, liver, heart, intestine, and bladder were removed and examined for helminths. All helminths from the first collection were heat fixed, stored in 70% ethanol, stained in acetocarmine, and mounted in Permount. Parasites from all collections were identified with keys and primary literature (Khalil et al., 1994; Harris and Vogelbein, 2006; Bray et al., 2008; Anderson et al., 2009).

### Statistical Analyses

To visualize the variation in parasite component community structure, we performed non-metric multi-dimensional scaling (NMDS). In the NMDS, average abundances of parasite species in each component community were calculated and square root transformed to control for the effect of high abundance parasite taxa. A resemblance matrix was constructed using Bray-Curtis dissimilarity distance, and stress was calculated using Kruskal’s stress formula 1. To determine whether there were significant differences in parasite component community structure, we performed a permutational multivariate ANOVA (PerMANOVA). For this analysis, we used site as a factor and performed 999 permutations. Additionally, we performed a 2-way crossed similarity percentage (SIMPER) analysis to identify the parasite species that contributed to 5% or greater of the variation in parasite community structure between sites (PRIMER-E LTD., Plymouth, U.K.).

To determine the role of landscape composition and physicochemical factors in determining parasite community composition, we used a mulitivariate regression tree using R packages rpart and mvpart (R Core Development Team, 2016; De’ath 2004; Therneau et al. 2019). Regression tree analysis is beneficial for determining the effects of urbanization on a response variable because the method does not assume independence between samples, and makes no *a priori* assumptions on the relationship between the response and predictor variables. Prior to this analysis, we conducted a variable reduction process to simplify the 48 water and 65 sediment trace metal and nutrient concentration data (JMP Pro 12.0.0, SAS Institute Inc). To do this, we generated a correlation matrix of element concentrations, placed all variables in one cluster, and then conducted binary splits based on the magnitude of the second eigenvalues. Variables were iteratively reassigned to clusters, and cluster membership was finalized when the squared correlation to the first principal component was highest. In each cluster, the variable that explained the most variation was used in the subsequent regression tree analysis. In the regression tree analysis, the multivariate response variable was the average abundances of the parasite species and the predictor variables were the landscape metrics and representative physicochemical factors.

## Results

### Parasite community in *Fundulus heteroclitus*

Eight taxa of metazoan parasites were identified from 630 fish, representing 24 different parasite communities. The direct life-cycle parasites included the branchiuran *Argulus funduli*, the copepod *Ergasilus funduli*, the leech *Myzobdella lugubris*, and the monogeneans *Swingleus ancistrus* and *Fundulotrema prolongis*. The indirect life-cycle parasites included the digenean *Lasiotocus minutus*, the cestode *Glossocercus caribaensis*, and the nematode genera *Contracaecum* sp. These 8 taxa of parasites infected more than 70% of the killifish examined (Tables 1 and 2).

**Table 1.**
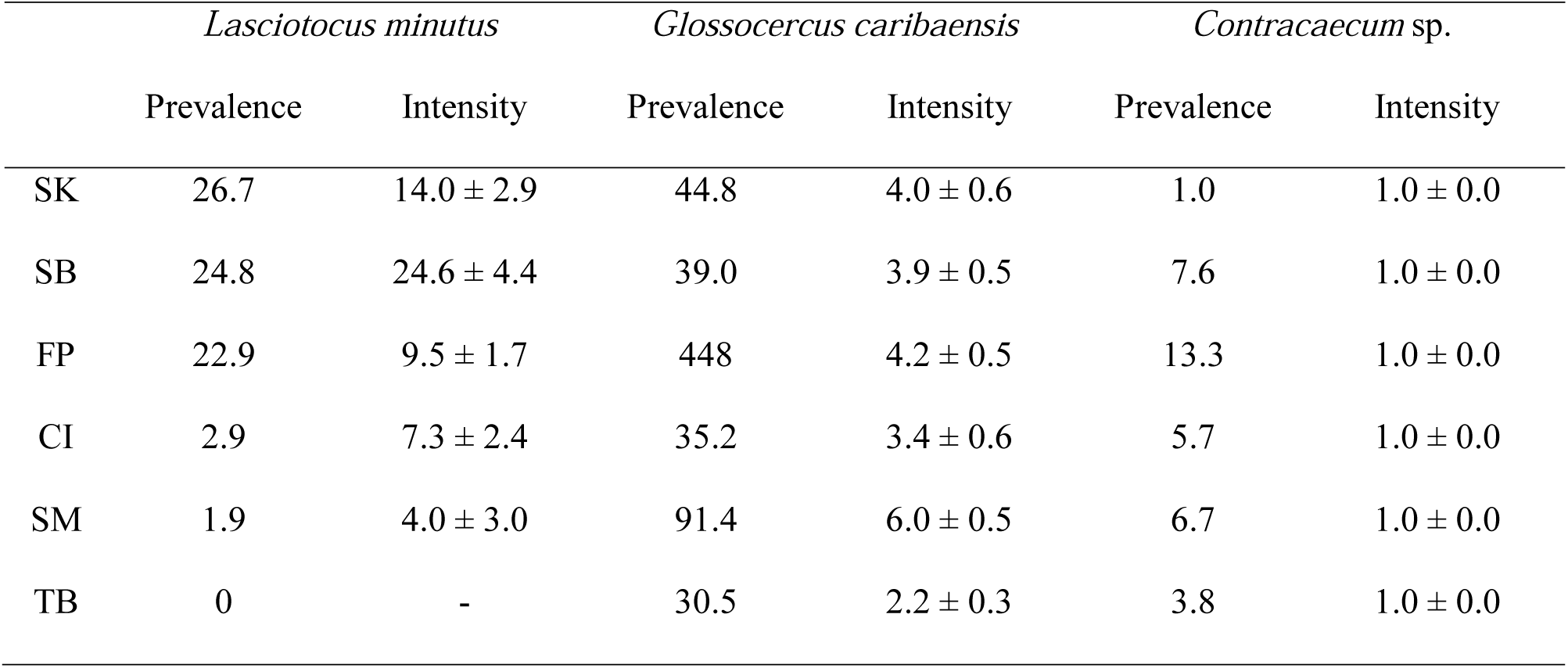
Prevalence (%) and mean intensity (SE) of the 3 complex life cycle parasite taxa infecting the salt marsh fish, *Fundulus heteroclitus*. The sites include: Fort Pulaski (FP), Cockspur Island (CI), Tybee (TB), and Skidaway Island (SK) in the Savannah River Estuary; Shellman Bluff (SB) as a representative of the Sapelo Sound; and St. Marys (SM) as part of the St. Marys Estuary. The sites are presented from top to bottom to reflect increasing levels of impervious surface.

| *Lasiotocus minutus* |  | *Glossocercus caribaensis* |  | *Contracaecum* sp. |  |
| --- | --- | --- | --- | --- | --- |
| Prevalence | Intensity | Prevalence | Intensity | Prevalence | Intensity |

|  |  |  |  |  |  |  |
| --- | --- | --- | --- | --- | --- | --- |
| SK | 26.7 | 14.0 ± 2.9 | 44.8 | 4.0 ± 0.6 | 1.0 | 1.0 ± 0.0 |
| SB | 24.8 | 24.6 ± 4.4 | 39.0 | 3.9 ± 0.5 | 7.6 | 1.0 ± 0.0 |
| FP | 22.9 | 9.5 ± 1.7 | 448 | 4.2 ± 0.5 | 13.3 | 1.0 ± 0.0 |
| CI | 2.9 | 7.3 ± 2.4 | 35.2 | 3.4 ± 0.6 | 5.7 | 1.0 ± 0.0 |
| SM | 1.9 | 4.0 ± 3.0 | 91.4 | 6.0 ± 0.5 | 6.7 | 1.0 ± 0.0 |
| TB | 0 | - | 30.5 | 2.2 ± 0.3 | 3.8 | 1.0 ± 0.0 |

**Table 2.** Prevalence and mean intensity (SE) of the 5 direct life cycle parasite taxa infecting the salt marsh fish, *Fundulus heteroclitus*. The sites include: Fort Pulaski (FP), Cockspur Island (CI), Tybee (TB), and Skidaway Island (SK) in the Savannah River Estuary; Shellman Bluff (SB) as a representative of the Sapelo Sound; and St. Marys (SM) as part of the St. Marys Estuary. The sites are presented from top to bottom to reflect increasing levels of impervious surface.

|  |  |  |  |  |
| --- | --- | --- | --- | --- |
| *Myzobdella lugubris* | *Swingleus ancistrus* | *Fundulotrema*<br>*prolongis* | *Ergasilus funduli* | *Argulus funduli* |

|  | Prevalence | Intensity | Prevalence | Intensity | Prevalence | Intensity | Prevalence | Intensity | Prevalence | Intensity |
| --- | --- | --- | --- | --- | --- | --- | --- | --- | --- | --- |
| SK | 0 | - | 40.0 | 2.8 ± 0.4 | 29.5 | 4.6 ± 1.0 | 24.8 | 3.4 ± 0.7 | 15.2 | 1.4 ± 0.2 |
| SB | 1.9 | 1.0 ± 0 | 36.2 | 3.1 ± 0.3 | 20.0 | 4.2 ± 0.9 | 19.0 | 3.8 ± 0.7 | 20.0 | 1.4 ± 0.2 |
| FP | 9.5 | 1.2 ± 0.2 | 38.1 | 4.5 ± 0.7 | 29.5 | 6.0 ± 1.1 | 14.3 | 3.6 ± 0.7 | 24.8 | 1.4 ± 0.1 |
| CI | 0 | - | 8.6 | 2.0 ± 0.5 | 16.2 | 3.6 ± 0.8 | 20.0 | 3.9 ± 1.1 | 7.6 | 1.1 ± 0.2 |
| SM | 0 | - | 13.3 | 2.7 ± 0.5 | 20.0 | 3.6 ± 0.9 | 22.9 | 3.4 ± 0.9 | 12.4 | 1.2 ± 0.2 |
| TB | 0 | - | 16.2 | 1.4 ± 0.3 | 19.0 | 3.5 ± 0.5 | 11.4 | 3.1 ± 0.5 | 10.5 | 1.3 ± 0.1 |

Assessed with NMDS and PerMANOVA, the parasite community composition differed between sites (*P* < 0.001, Pseudo F_5,12_ = 6.11; Fig 2). SIMPER analysis indicated that the observed differences in parasite community structure were driven by indirect life-cycle parasites: the digenean *Lasciotocus minutus*, and the cestode *Glossocercus caribaensis*. This analysis demonstrated that direct life-cycle parasites did not contribute to the variation in parasite community structure.

**Fig 2.**
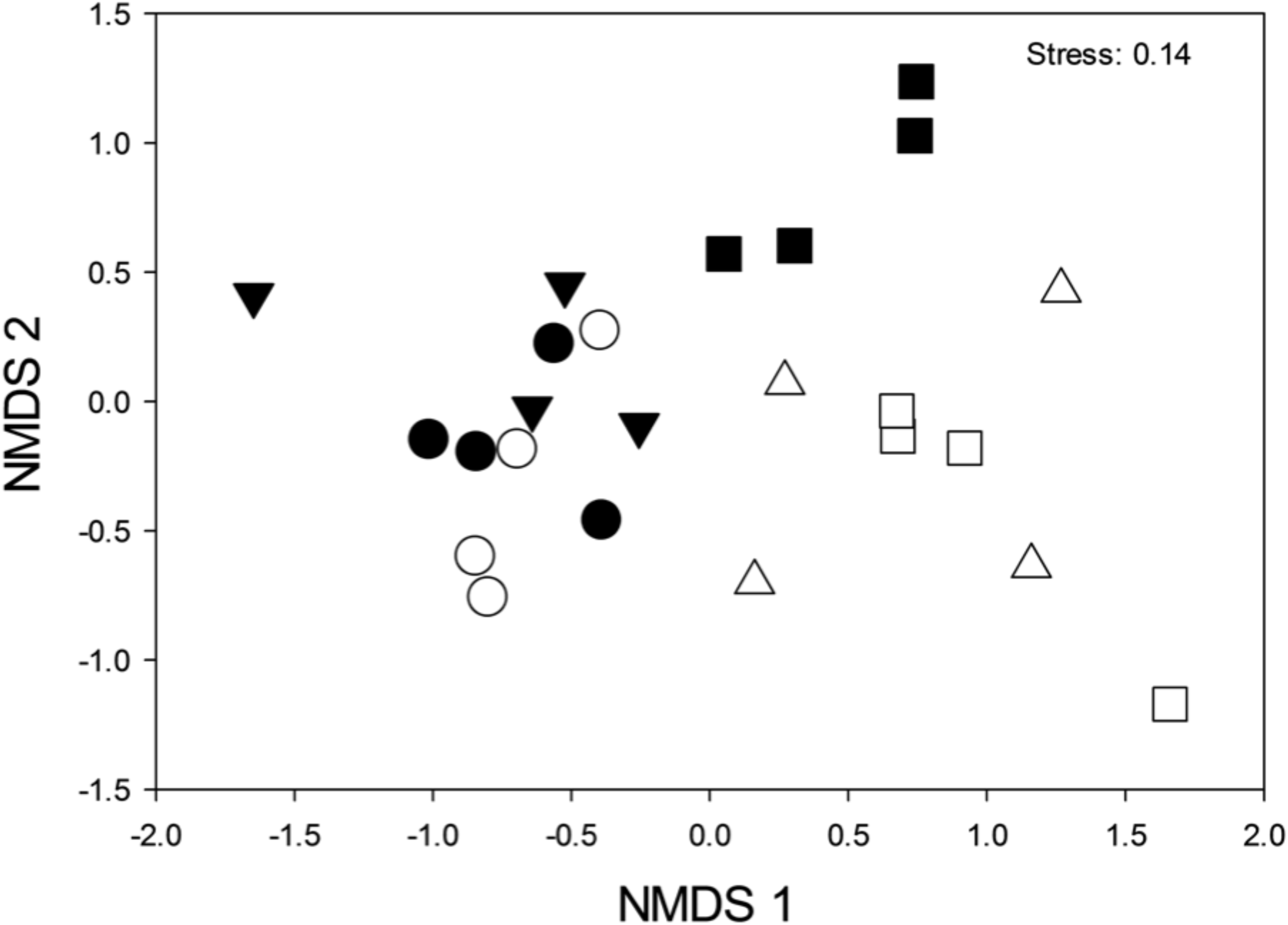
Non-metric multidimensional scaling plot of complex and direct life cycle parasite component communities of the salt marsh fish, *Fundulus heteroclitus*, from 6 sites in coastal Georgia, USA. Distances are based on Bray-Curtis dissimilarities of the square root transformed mean abundance of 8 parasite taxa of *Fundulus heteroclitus*. The sites include: Fort Pulaski (FP) (closed triangles), Cockspur Island (CI) (open triangles), Tybee (TB) (open squares), and Skidaway Island (SK) (open circles) in the Savannah River Estuary; Shellman Bluff (SB) (closed circles) as a representative of the Sapelo Sound; and St. Marys (SM) (closed squares) as part of the St. Marys Estuary.

### Landscape and physicochemical predictors of parasite community in *Fundulus heteroclitus*

The cluster analysis variable reduction process on the water and sediment trace metal and nutrient concentrations resulted in 16 clusters, explaining 83.3% of the variance. We constructed a multivariate regression tree that designated each of the 24 parasite community observations (6 sites, 4 collection periods) to terminal leaf nodes: these terminal leaf nodes categorize the factors, where the factors are all the collected landscape and physicochemical variables collected from the 6 sites across 4 collection periods that predict the observed parasite community structure. In multivariate regression tree analysis, variable importance is determined as the reduction in the loss function attributed to each variable at each split summed across the full tree. The optimum tree size contained 3 leaves, and the primary variable that was most important in determining the parasite community structure was the potassium concentrations in the sediment, with largest patch index (LPI) and aquatic osmium concentrations as secondary predictor variables (CV Error = 0.5608). Parasite communities that were dominated by the indirect life-cycle cestode *Glossocercus caribaensis* were characterized by sediment potassium levels with a mean of 4063 +/-270 mg/L, and an LPI of less than 22. Parasite communities that were dominated by the indirect life-cycle digenean *Lasiotocus minutus* were characterized by sediment potassium levels with a mean of 4706 +/-468 mg/L and an osmium concentration of greater than 0.01 mg/L.

## Discussion

The goal of this study was to determine the effects of urbanization, mediated through alterations in landscape composition and physicochemical factors, in structuring parasite communities. We determined that parasite community composition differed between sites with different levels of urbanization. Variation in the prevalence and intensity of infection by indirect life-cycle parasites, the digenean *Lasiotocus minutus* and the cestode *Glossocercus caribaensis* were the primary drivers for the differences in parasite communities. Although the prevalence of direct life-cycle parasites differed among sites, this difference did not contribute to the overall variation in community composition. Additionally, we found that the landscape composition and physicochemical factors that best predicted the parasite communities were potassium sediment concentration, largest patch index, and aquatic osmium concentration. These data support the prediction that in the common marsh killifish, parasite community composition is explained by metrics associated with human modification of the environment, and parasite responses to urbanization are dependent upon life history.

Our results indicated that sediment potassium, water osmium, and largest patch index were the most important factors in determining the structure of parasite communities. Although potassium is not necessarily a contaminant, the concentrations we observed have been reported to significantly affect estuarine communities (Ajitthamol et al. 2016). The upstream watersheds of the high potassium sites are dominated by cotton and soy agriculture, which are especially dependent on potassium fertilizers. The other major contaminant we found to structure parasite communities is the rare element osmium. The concentrations that we observed suggest extreme point source pollution (Turekian et al. 2007). Taken together, these data suggest that our sites may have contamination that impacts the survival of benthic organisms, which may subsequently alter trophic interactions and parasite transmission that relies upon those interactions. The major landscape factor in determining parasite community composition, LPI, is a simple measure of landscape “dominance.” LPI typically decreases as distance from urban centers increase (Nong et al. 2018). We found an LPI value of 22 to best predict parasite community structure, and this value was also associated with lower sediment and aquatic contaminant levels. This metric suggests that urbanization is a major driver of parasite community variation, whereby areas with lower levels of urbanization likely support more diverse and stable community of organisms that allow the completion of indirect parasite life-cycles.

Previous studies have shown a relationship between urbanization and parasite community variation. Blanar et al. (2016) demonstrated a correlation between parasite community structure and local contaminants and land use within 5 km. In this study, the relative abundance of the indirect life-cycle digenean was positively associated with an increase in crude oil contaminants, whereas the opposite pattern was observed for the directly transmitted monogenean parasites. This observation follows results of an earlier meta-analysis, where parasite life history (>1 obligatory host in life-cycle vs. 1 obligatory host in life-cycle) and habitat (external vs. internal) informed whether aquatic pollution had a significant effect on parasite population dynamics (Blanar et al., 2009). Additionally, Calegaro-Marques and Amato (2014) revealed an association between an urban-rural gradient and life-cycle variation in parasite communities of Rufous-bellied thrushes. Our results concur with these studies, and support the proposition that urbanization disproportionately affects indirect life-cycle parasites, which is then reflected in the parasite community structure.

We found that sites within urbanized watersheds contained a greater prevalence and intensity of the autogenic (whole life-cycle is aquatic) parasite *Lasiotocus minutus*, a digenean that uses the focal fish species, *Fundulus heteroclitus*, as the definitive host, molluscs as the first intermediate host, and fish or crustaceans as the second intermediate host (Stunkard, 1980). This pattern was unexpected, as this parasite and associated hosts are continuously exposed to the external aquatic environment, and would be exposed to any contaminants or deleterious physicochemical parameters. However, the most important physicochemical parameter in structuring the parasite communities, sediment potassium, may alter trophic structures in a favorable way to the host communities. For example, potassium has been shown to increase phytoplankton abundance in estuaries. This increase in basal resource would disproportionately affect benthic grazers, such as the 1st intermediate hosts for *L. minutus*. The second complex life-cycle parasite, the cestode *G. caribaensis*, is allogenic (life-cycle is aquatic and terrestrial), and did not follow the same pattern as *L. minutus*, and was more dominant in sites associated with less impacts of urbanization. This allogenic cestode has fish-eating birds as definitive hosts and infects a number of fish intermediate hosts (Scholz et al., 2004). This parasite may appear to be sensitive to urbanization because its definitive host is vagile and can select habitat outside of urbanized salt marshes. Consequently, the difference in prevalence across sites may be due to the effect of the abiotic conditions on parasite life stages external to the host, or through effects on critical aquatic life-cycle hosts, such as the mollusc and bird (Anderson and Sukhdeo, 2010). Future research should include population size assessments of intermediate and definitive hosts to better disentangle the direct effects of urbanization (through exposure to the environment) and indirect effects (through population dynamics of the hosts).

Coastal urbanization and anthropogenic disturbance are rapidly altering free-living and parasite community dynamics. Our results suggest that urbanization and its effects are important in structuring parasite communities of the common marsh killifish, complex life-cycle parasites are more affected than direct life-cycle parasites, and the degree to which individual parasite species respond to urbanization is determined by their specific life history strategy. Further, our data suggest that urbanization likely affects interactions of free-living hosts, and their environment which is then detected as variation in the parasite community.

## Acknowledgements

We acknowledge the advice and support provided by Dr. Clark Alexander and Mike Robinson at the Applied Coastal Research Laboratory (Georgia Southern University), and Dr. Sayed Hassad at the Laboratory of Environmental Analysis (University of Georgia). Field assistance was provided by Emily Dodd, Sarah Dunn, Maria St. Jean, and Jackson Tomlinson. Comments on an earlier draft by Drs. Risa Cohen, James Roberts, and Oscar Pung improved this manuscript. This research was funded in part by a Research Assistantship from the Institute of Coastal Plain Science (Georgia Southern University) to JMA. TKA was supported by the Office of Research Services and Sponsored Programs at Georgia Southern University (GSU) and by an appointment to the USDA-ARS Research Participation Program administered by the Oak Ridge Institute for Science and Education (ORISE) through an interagency agreement between the U.S. Department of Energy (DOE) and USDA under contract number DE-AC05-06OR23100. The funders had no role in study design, data collection and interpretation, or the decision to submit the work for publication. Mention of trade names or commercial products in this article is solely for the purpose of providing specific information and does not imply recommendation or endorsement by the USDA. USDA is an equal opportunity provider and employer.

## Notes

#### Summary of Updates

Introduction revised, statistical analyses updated, and discussion modified.

